# Superior batch alignment and hyper-dimensional cytometry representations allow ultra-sensitive classification of disease phenotypes

**DOI:** 10.1101/2025.07.28.666458

**Authors:** Benjamin S. Mashford, Timothy Hewitt, Maryam May, Zixin Zhuang, Akshat Jain, Koula E. M. Diamand, Fei-Ju Li, Kristy Kwong, Stuart H. Read, Ainsley R. Davies, Dillon Hammill, T. Daniel Andrews

**Affiliations:** - The John Curtin School of Medical Research, The Australian National University, Canberra, Australia; – Computational Science Cluster, School of Computing, College of Systems and Society, The Australian National University, Canberra, Australia; – Australian Phenomics Facility, The Australian National University, Canberra, Australia

**Keywords:** cytometry, machine learning, bioinformatics

## Abstract

Analysis of cytometry data predominantly relies on clustering and dimensionality reduction approaches for computational tractability. This is particularly relevant for modern spectral flow cytometers, which can simultaneously measure an increasingly large number of antibody marker channels. While dimensionality reduction provides for more efficient data processing, this comes at the expense of data loss that may miss subtle patterns among rare cell types that may be critical for disease detection. Maintaining analysis at full dimensions presents opportunities to preserve resolution and provides complete downstream data interpretability. However, a significant obstacle to analysis of cytometry data without dimensionality reduction is the significant batch effects observed in this data between experiment days, operators and equipment types. Here we show a new strategy to denoise batch variation from both flow- and mass-cytometry datasets using an autoencoder neural network architecture. We generated a benchmark flow cytometry dataset in mice to compare this approach to current toolsets and find our approach shows superior preservation of biological signals, whilst also performing batch correction equal to current best methodology. Our batch alignment approach works to such an extent that it becomes practically possible to project batch aligned data into multidimensional space to generate a novel representation of cellular phenotype for downstream model building. This hyperdimensional approach maintains original data resolution without requiring dimensionality reduction, and thus any resultant cell populations that differentiate phenotypes remain fully interpretable. We show with two large clinical datasets that our batch-alignment approach coupled with the multi-dimensional representation successfully detects meaningful patterns in cases where the original analysis methods struggled. This new framework removes some of the inherent technical limitations encountered in the integration of large, multi-batch cytometry datasets and provides a framework for machine learning model building from this modality. An implementation of this framework and an associated web application accompanies this manuscript at http://voxelcoder.cloud.

**Significance Statement:** This work addresses a critical bottleneck in cytometry analysis by introducing a neural network-based approach that effectively removes technical batch variation while preserving biological signals. The novel hyperdimensional representation maintains full data resolution without dimensionality reduction, enabling more sensitive detection of disease-associated cellular signatures than current methods. Importantly, this framework enables reliable batch normalization of fresh samples processed at different times and locations, overcoming the practical constraints of clinical sample collection where simultaneous processing is often impossible. The enhanced sensitivity for identifying pathogenic cellular patterns has immediate implications for biomarker discovery and personalized medicine applications.

## Introduction

Quantitative analysis of the cellular phenotype is routinely performed in a research and clinical setting through simultaneous measurement of cellular markers that target surface proteins, nucleic acids and other molecular components. Common modalities of this routine cellular assay include both flow cytometry^1,2^ and CyTOF mass cytometry^3^. Two technical challenges limit the automated interpretation of these data: batch effects in data acquisition and the complexity of analyzing high-dimensional marker combinations^4^. To counter these difficulties, current methodologies rely predominantly on manual gating, which, while established for standardized panels, becomes increasingly challenging as the number of measured parameters increases^5^, as is now routinely possible with spectral flow cytometry^6^.

Batch effects in flow cytometry arise from multiple technical sources during data acquisition. These include variations in reagent preparation, such as differences in antibody clone selection and fluorophore conjugation between batches, as well as variations in reagent concentrations that affect staining intensity. Operator-dependent factors introduce additional variability through differences in sample preparation technique, instrument setup, and data acquisition protocols. Furthermore, inherent drift in instrument calibration over time affects both laser intensity and detector sensitivity, leading to systematic shifts in fluorescence measurements. Despite implementation of standardized protocols and quality controls^7,8^, these technical variations between experimental batches persist, manifesting as shifts in marker intensity distributions. These variations present particular challenges for data integration across multiple studies or institutions^7^, especially in multi-centre studies with requirement to pool samples to improve overall study power. While computational approaches have demonstrated potential for addressing batch effects^9–14^, much scope still exists for practical improvement of these methodologies ^4,5,15–17^.

The increasing number of parameters measured by modern flow cytometers introduces additional analytical complexities^6^. Standard manual gating in two dimensions requires examination of numerous scatter plots potentially obscuring subtle correlations between markers^7^. For instance, a six-marker panel necessitates analysis of 15 distinct two-dimensional plots, and this quickly increases with addition of extra markers, becoming unfeasible for exhaustive visual analysis with the marker numbers now possible with spectral flow cytometry. Gating strategies to identify pre-defined cell populations of interest partially address this difficulty^18^, yet this also works to obscure unexpected changes. Dimensionality reduction techniques have been applied to facilitate visualization and analysis of high-dimensional cytometry data^9^. However, such approaches necessarily reduce data resolution, potentially obscuring rare cell populations that may have biological significance ^16^.

Recent computational methods have proposed alternative strategies for analysing high-dimensional cytometry data. The adaptation of single-cell RNA sequencing analytical techniques to flow cytometry data^19^ suggests new approaches to population identification. Additionally, investigations into hyperdimensional computing have demonstrated the feasibility of analysing flow cytometry data without dimensionality reduction^20^. Analysis of multi-parametric flow- and-mass cytometry datasets in hyper-voxel space conceptualises each individual cell as a data point in a multi-dimensional lattice. Each voxel, or hypercube, is a discrete region of the multi-dimensional parameter space, where each dimension is defined by variation of a single marker intensity. Hyper-dimensional computing frameworks can efficiently encode and manipulate high-dimensional representations, facilitating the extraction of meaningful insights from noisy and heterogeneous data sources^21^. One example of alignment of biological manifolds containing cytometry data employed a generative adversarial network^22^. While further work in this area has used quadratic form cluster matching to allow multi-dimensional batch alignment^23^.

We present a novel computational toolset and an implementation (including a web tool at http://voxelcoder.cloud) that combines deep learning-based batch alignment with systematic multi-dimensional cell population analysis. This employs an autoencoder^19^ neural network architecture to perform batch correction against a reference distribution, effectively normalizing technical variations while preserving biological signals. Following batch alignment, the pipeline implements a novel “voxel-gating” strategy that systematically analyzes cellular populations across multiple marker combinations without requiring dimensionality reduction. This is achieved by dividing each marker’s expression range into discrete intervals and examining all possible marker combinations to generates comprehensive, unbiased representations of cellular phenotypes that remain fully interpretable. The resulting discretized population signatures can be used to build robust classification models for diagnostic applications. An implementation of this novel framework, named VoxelCoder, accompanies this work. We demonstrate that VoxelCoder outperforms traditional manual gating approaches while eliminating the need for per-plate normal controls and manual intervention. We evaluate VoxelCoder on a number of multicolor flow cytometry and CyTOF mass cytometry datasets and demonstrate classification performance while maintaining interpretability of results.

## Results

### Batch alignment

We developed a new approach to integration of cytometry data that employs an autoencoder neural network architecture to perform batch correction (Figure 1). We employ this architecture using a randomly chosen reference batch of samples collected at a single timepoint to establish the target distribution. The autoencoder learns to transform input samples to match the reference batch distribution while preserving biological variation. The autoencoder is implemented with batch normalization layers and trained using both reconstruction loss and distribution matching via histogram loss. Briefly, the network consisted of three encoding layers (64, 32, and 16 nodes) and three symmetrical decoding layers, with Rectified Linear Unit (ReLU) activation functions between layers. The autoencoder was trained to transform input data from different batches to match a reference distribution. For a given multi-batch dataset, one batch is randomly selected as the reference, and a model is trained using a combination of mean squared error (MSE) loss and a custom histogram loss function that encouraged the transformed data to maintain similar marker distribution shapes as the reference batch. The histogram loss term is computed by comparing normalized histogram distributions between the network output and reference data across all markers, using Wasserstein distance as the similarity metric. This dual loss function helps ensure that both individual cell measurements and overall population distributions are preserved during batch alignment.

**Figure 1.**
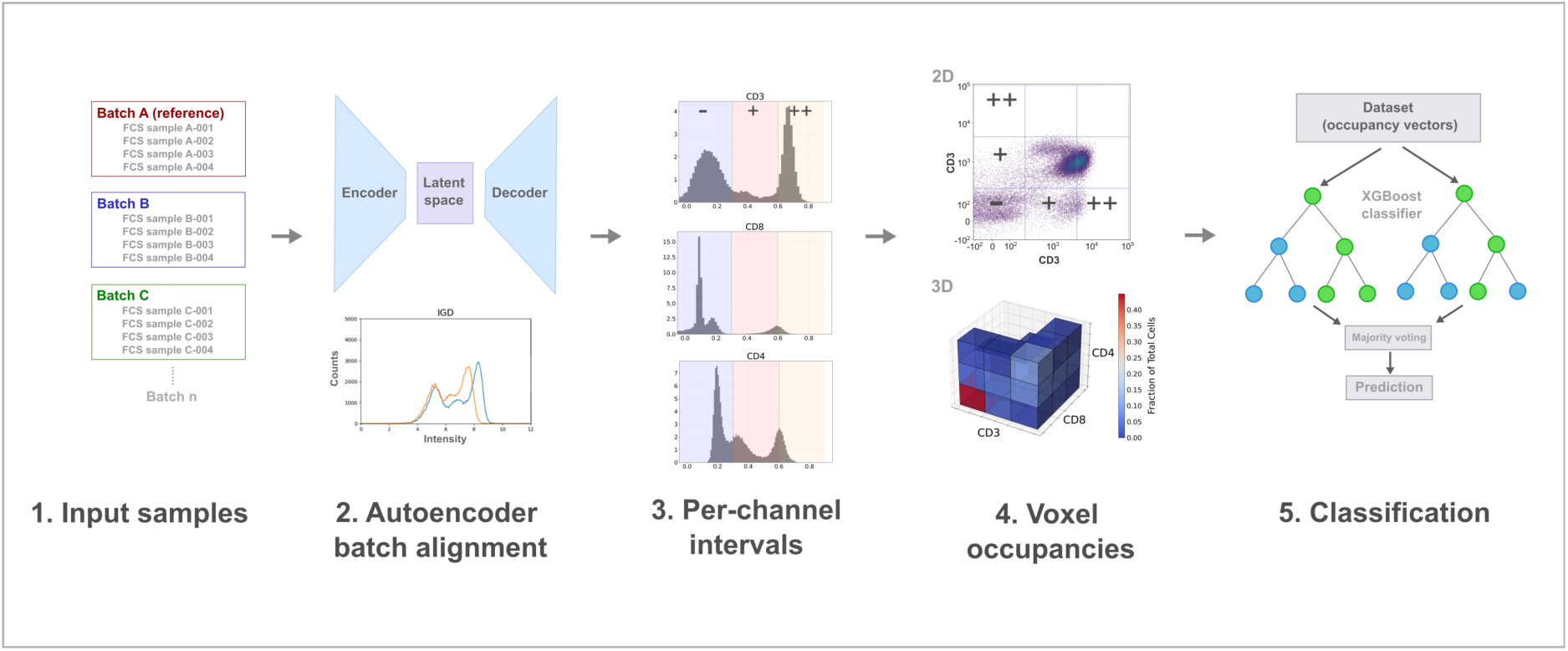
Overview of computational pipeline for automated flow cytometry analysis, beginning with input sample batches, proceeding through autoencoder-based batch alignment and application of fixed-size per-channel intervals, followed by voxel occupancy calculation, and machine learning classification.

### Voxel gating

Following batch alignment, we introduced a systematic *hyper-voxel gating* analysis strategy to comprehensively characterize cell populations across multiple markers. This method divides each marker’s expression range into three intervals (low, medium and high; denoted -, + and ++, respectively), based on the density distribution of the reference batch, and then exhaustively analyzes all possible 2,3,4 marker combinations. For each combination, we compute cell occupancy frequencies across the resulting n-dimensional voxel space (e.g. the three intervals generates 3 × 3 × 3 = 27 voxels per marker combination in 3D), generating a high-dimensional feature vector that captures detailed information about cell population distributions.

While conventional analysis of flow cytometry data relies on manual gating strategies where cell populations are sequentially identified using two-dimensional projections of the data, such approaches are limited to examining predefined regions of interest and may not capture cell populations that exist in unexplored marker combinations. The hyper-voxel gating methodology described here addresses these limitations while maintaining interpretability, like manual gating. The resulting hyper-voxel occupancy features serve as input for subsequent machine learning classification tasks, providing a comprehensive and unbiased representation of cellular phenotypes that may include both previously identified and potentially novel cell populations.

### Benchmarking of batch alignment

We evaluate our batch-alignment and voxel-gating approach with one purpose-generated benchmark dataset and two real-world datasets. The benchmark dataset is a biologically-replicated, multicolour flow cytometry experiment of C57/BL6 mouse spleen cells with a synthetic batch effect introduced between technical replicates by varying the dilutions of the antibodies (see Methods). In addition to this, the real-world data are two public datasets, including i) mass cytometry data of latent cytomegalovirus infection^24^, and ii) a multi-modal dataset of both flow- and mass-cytometry from individuals infected with COVID-19 and their clinical outcome^25,26^. In evaluating performance of batch-alignment of these data, the hyper-voxel gating strategy described above provides a ready framework for benchmarking.

### Alignment of synthetic batch effect

As mentioned, to appraise the performance of batch alignments, we generated a specific benchmark dataset with splenic cells from three wild-type C57BL/6 mice (see Methods). These wild-type mice represent biological replicates, and for each of these a blood sample was analysed in three technical replicates, where a synthetic batch effect was introduced by varying the concentration of antibodies included in the marker panel used for each of the three replicates (see Methods, Figure 2 & Table 4). The panel included 11 markers: B220, CD3, CD8, CD19, CD25, CD44, CD62L, IgD, IgM, Ly6C, NK1 and a viability dye. Prior to analysis, data underwent standard pre-processing including compensation and debris removal. In this benchmark dataset, the technical replicates are derived from the same spleen cell draw from the same individual mouse. A perfect batch alignment of the flow cytometry information from the three replicates should result in a near-identical distribution of marker intensities and cell counts in hyper-voxel.

**Figure 2.**
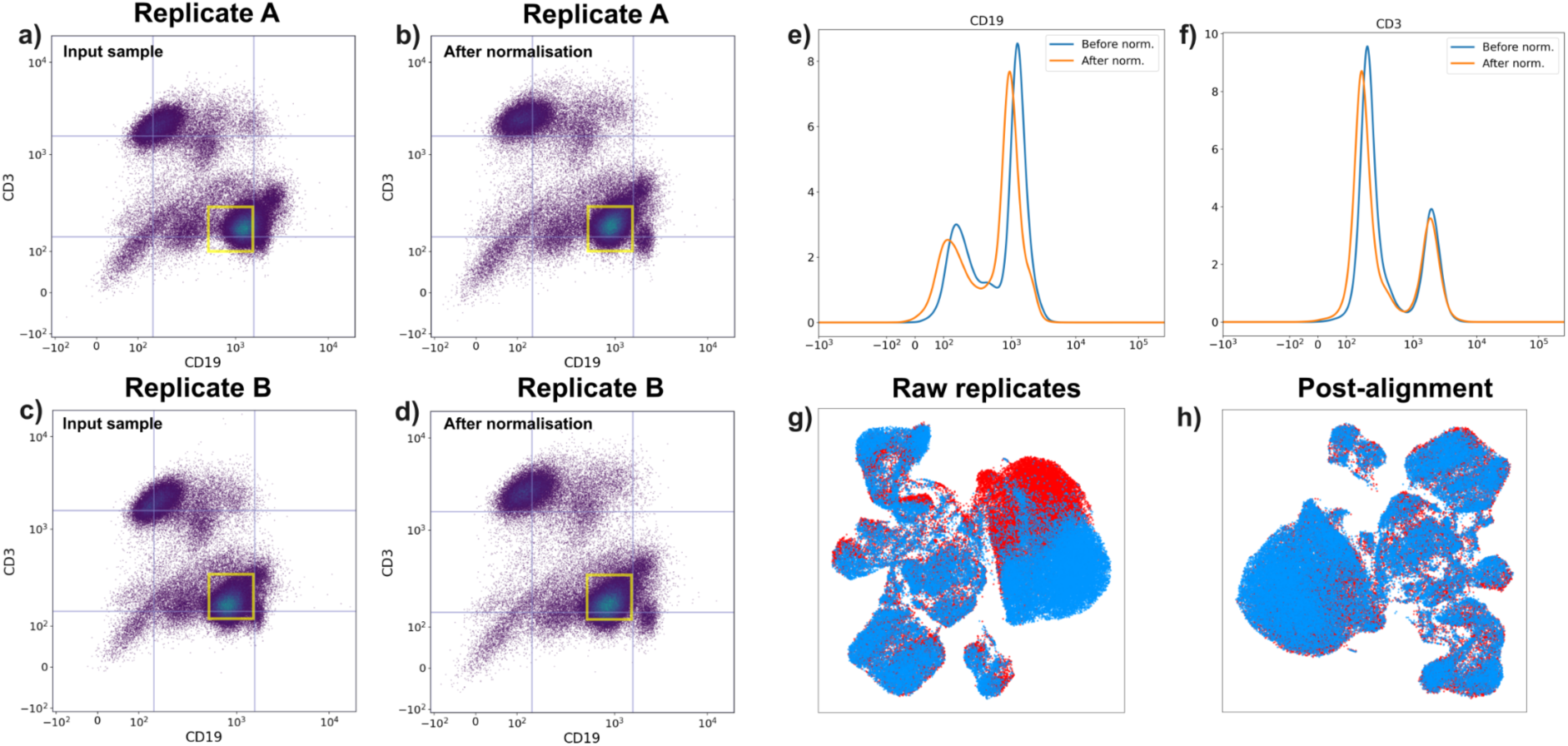
Batch alignment with an autoencoder neural network architecture. a-d) Two-dimensional scatter plots of CD3 and CD19 marker intensities for two technical replicates that harbour an inter-replicate synthetic batch effect introduced by varying antibody marker concentration. Replicate A (a & b) and Replicate B are aligned using an autoencoder model, as implemented in VoxelCoder. The squares in yellow (a-d) are in a constant position between plots and shows the movement of the cells with batch alignment (between the input sample and the normalised output). The scatterplot color scale is proportional to cell density. e-f) Histograms showing removal of batch-related shifts in signal intensities for two fluorescence channels for the CD19 and CD3 markers from the same replicate shown in a and b. The blue lines indicate the distribution of intensities prior to batch alignment and the red lines are after normalisation. g-h) UMAP visualization demonstrating batch alignment of two technical replicates (blue replicate superimposed over red replicate) with introduced synthetic batch effects. The raw replicate signals show clear batch effects (g) and post-alignment samples exhibit improved overlap of cell populations (h).

Figure 2a-d shows the effectiveness of our autoencoder-based batch correction method across technical replicates of the same cell suspensions from three biological replicate mice. The technical replicates and the synthetically introduced batch effect they harbour is effectively removed by the autoencoder neural network described above. The input samples show clear batch-specific variations in both the CD3 and CD19 dimensions (Figure 2, a-d), particularly evident in the position and shape of the CD3^-^ CD19^+^ B cell population (lower right cluster with yellow box). The autoencoder successfully normalized this introduced batch variation while preserving the biological distinctions between cell populations. This preservation of the structure of cell populations is important, as it indicates the method can distinguish between technical artifacts and true biological variation. The corresponding histogram analyses (Figure 2e-d) provide quantitative evidence of the normalization, showing how the method aligns both the location and shape of marker distributions to the reference batch. Notably, the correction handles non-linear distortions in signal distributions, as seen in the varying peak positions and shapes of the input histograms, suggesting the method is robust to complex batch effects. The aligned outputs maintain consistent population boundaries across all three batches while preserving the relative proportions of major cell populations, demonstrating that the autoencoder learns a transformation that removes technical variation without compromising biological signal. Shown in Figure 2g, prior to batch correction, cells from the same biological sample processed in different batches form non-overlapping clusters. After autoencoder-based alignment (Figure 2h), these cells merge, demonstrating successful normalisation of batch effects while preserving the underlying cell population structures.

### Comparison with existing toolsets

Additionally, we benchmarked this autoencoder neural network approach, implemented by VoxelCoder, against three popular batch alignment methods: CytoNorm, CyCombine, and Harmony. Supplementary Figure 1 shows comparison of two-dimensional scatterplots pre- and post-batch alignment. To benchmark these tools, we use two differing benchmarking approaches. The first is through comparison of the similarity of the distributions of intensities for each marker channel. For this we used per-channel Wasserstein Distances (Table 1). Complementary to this, we secondly employed hypervoxel-based Euclidean distances (Table 2). These complementary approaches provide a comprehensive assessment of batch correction performance, with per-channel analysis focusing on individual marker distributions and hypervoxel analysis capturing high-dimensional phenotypic relationships.

**Table 1.** Wasserstein distances between technical replicate mice after batch alignment with different methods.

| Sample | Non-aligned | VoxelCoder | CytoNorm | CyCombine | Harmony |
| --- | --- | --- | --- | --- | --- |
| Mouse 1 (male) | 0.080 | 0.081 | 0.085 | 0.060 | 0.055 |
| Mouse 2 (female) | 0.085 | 0.030 | 0.089 | 0.061 | 0.027 |
| Mouse 3 (female) | 0.052 | 0.024 | 0.061 | 0.036 | 0.028 |

**Table 2.** Euclidian distances calculated between technical replicate mice, using hypervoxel representation, after batch alignment with.

| Sample | Non-aligned | VoxelCoder | CytoNorm | CyCombine | Harmony |
| --- | --- | --- | --- | --- | --- |
| Mouse 1 (male) | 0.145 | 0.028 | 0.188 | 0.054 | 0.030 |
| Mouse 2 (female) | 0.180 | 0.043 | 0.288 | 0.060 | 0.032 |
| Mouse 3 (female) | 0.185 | 0.053 | 0.220 | 0.087 | 0.042 |

The removal of batch effects has two important components to consider. First, the distributions of signal values need to be brought to a similar mean and variance between batches to allow simplified direct comparison. However, secondly, this must be achieved whilst also preserving the biologically-relevant features that are the real signal differences between samples. For example, it is simple enough to fit any distribution present in an input dataset to a given mean and variance, yet this will likely erase the real signal present in the data. We compared the popular batch alignment methods with our autoencoder normalisation approach for both similarity of input distributions and preservation of biological signal following normalisation.

The similarity of normalised distributions from different popular tools using per-channel distributions reveals that Harmony generally achieves the lowest Wasserstein Distances across most mouse samples (Table 1). Wasserstein Distance is a metric that represents the subtraction of one distribution from another, and is colloquially known also as Earth Mover’s Distance. Lower Wasserstein Distance indicates effective alignment between technical replicates. VoxelCoder demonstrates competitive performance, particularly for certain samples, though it exhibits slightly higher distances for others. CyCombine also performs adequately with consistent distances across all samples examined. The hypervoxel analysis provides an assessment of batch effect correction methods by characterizing cellular phenotypes in high-dimensional space, offering a complementary perspective to the per-channel Wasserstein approach. Using hyper-voxel cell counts, we calculate the Euclidean Distance between input and normalised counts, using this multidimensional representation. Table 2 shows both VoxelCoder and Harmony demonstrate effective performance across the samples tested. Harmony achieves the lowest Euclidean distances for most samples, while VoxelCoder performs comparably well.

When considering retention of biological signal following batch normalization, we compared the inter-biological replicate differences of the wild type mouse and mutant mouse synthetic batch effect data (Figure 3). These synthetic benchmarks provide a unique validation of the technical performance of the autoencoder framework. To quantify biological signal retention, we calculated Kullback-Leibler (KL) divergence between all sample pairs, where larger divergences between biologically distinct samples (wild-type vs mutant strains) relative to technical replicates indicate better preservation of biological variation. The synthetic dataset design allows clear separation of biological from technical variation since expected biological differences are known a priori, creating an ideal testbed for evaluating batch correction methods. This analysis shows that the autoencoder normalisation implemented in VoxelCoder retains more biological signal than other popular tools, including Harmony, whilst achieving similar utility for alignment of marker intensity distributions.

**Figure 3.**
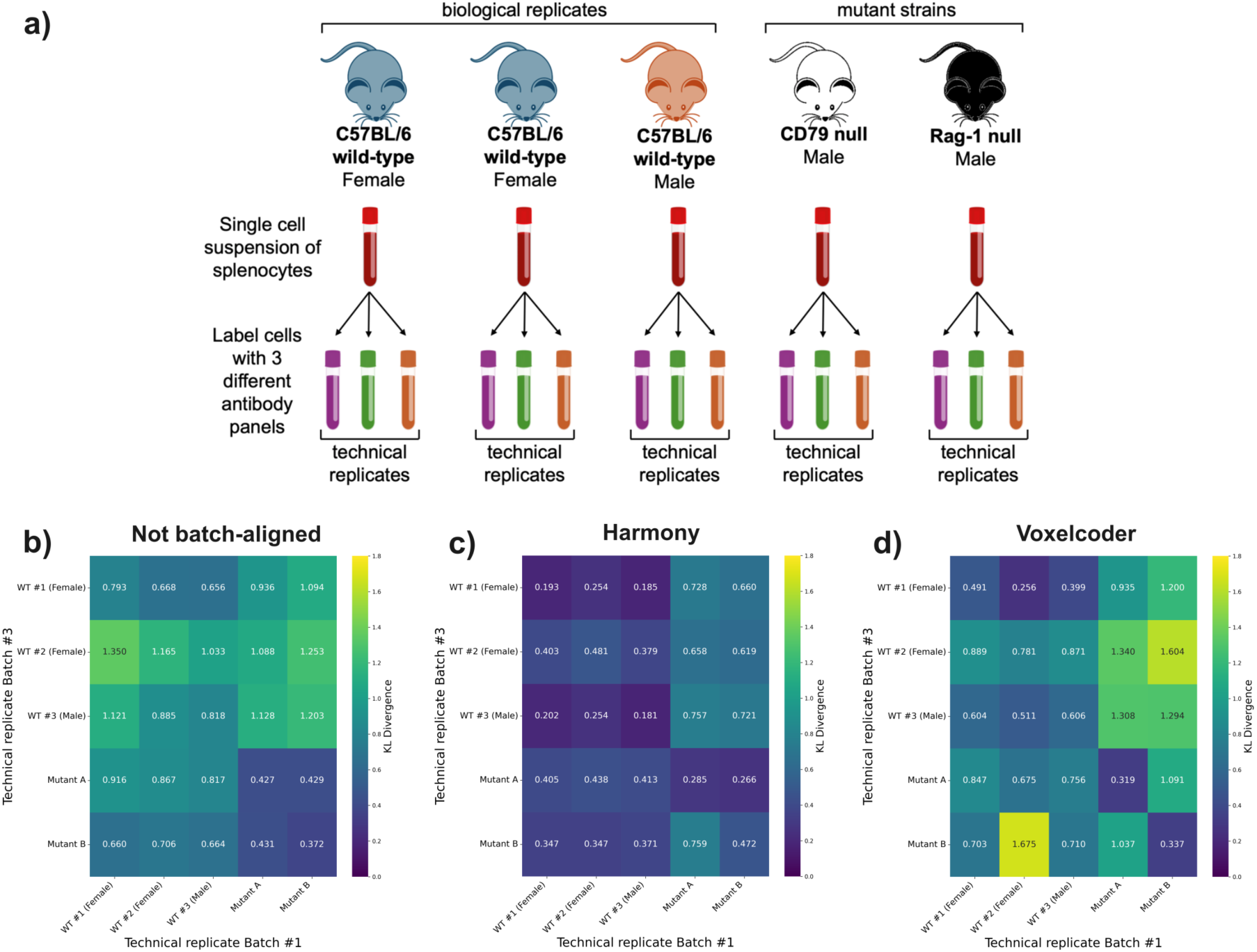
Upper row. **(a)** Schematic of design of synthetic batch effect mouse dataset. Spleens from three individual wild-type C67BL/6 mice (1 male, 2 female) provided cell samples for biological replication. A further two mutant C57BL/6 strains provided examples biological variation due to genetic changes. Each suspension of splenocytes from each mouse was split into three technical replicate vials of equivalent total cell count. These technical replicates for each mouse were prepared for fluorescence-activated cell sorting (FACS) with one of three marker cocktails. The marker cocktails contained identical antibodies with deliberate, minor differences to antibody dilutions (Table 4) to introduce a synthetic batch effect from the same input cells per biological replicate. **(Lower row)** Heatmap plots showing KL-divergence distances between 5 mice in the synthetic batch effect dataset. Dataset includes 3 wild-type mice (WT #1,2,3) and 2 mice with significant mutant phenotype (Mutant A, B). Scores are derived from per-channel histograms, using 50 bins per channel. The figures shows distances in non-batch aligned dataset **(b)**, compared to distances in Harmony-aligned **(c)** and VoxelCoder-aligned **(d)** datasets.

### Identification of Latent Cytomegalovirus Infection through Mass Cytometry Data Analysis

To assess our method against earlier work, using real-world data, we utilized a comprehensive cytometry by time-of-flight mass spectrometry (CyTOF) dataset^24^. This dataset comprises 472 samples from nine independent studies, containing peripheral blood mononuclear cell (PBMC) measurements from healthy individuals along with their cytomegalovirus (CMV) serostatus. The dataset represents a challenging real-world scenario due to its heterogeneous nature, combining data from multiple independent studies, making it particularly suitable for evaluating batch alignment analytical methods for cytometry data.

We conducted a re-analysis of this data in a hyper-voxel framework with VoxelCoder (Figure 4). Initial UMAP visualization (Figure 4a) clearly shows batch-specific clustering before alignment, with samples from each study forming distinct clusters. Following batch alignment with VoxelCoder (Figure 4b), we observe substantially improved mixing of samples across batches. Representative flow cytometry scatter plots (Figure 4c) demonstrate our voxel-gating approach, showing how intensity intervals for key markers (CD3, CD19, CD4, CD8, CD27, CD20) are partitioned into low (-), medium (+), and high (++) expression categories. These classifications form the basis for the hyper-voxel framework used in our subsequent analysis. The Receiver Operating Characteristic (ROC) curve (Figure 4d) shows that our VoxelCoder-based model achieves an area under the curve (AUC) of 0.90 for CMV status prediction, outperforming the Harmony-aligned voxel analysis which achieved an AUC of 0.82 (Supplementary Figure 3).

**Figure 4.**
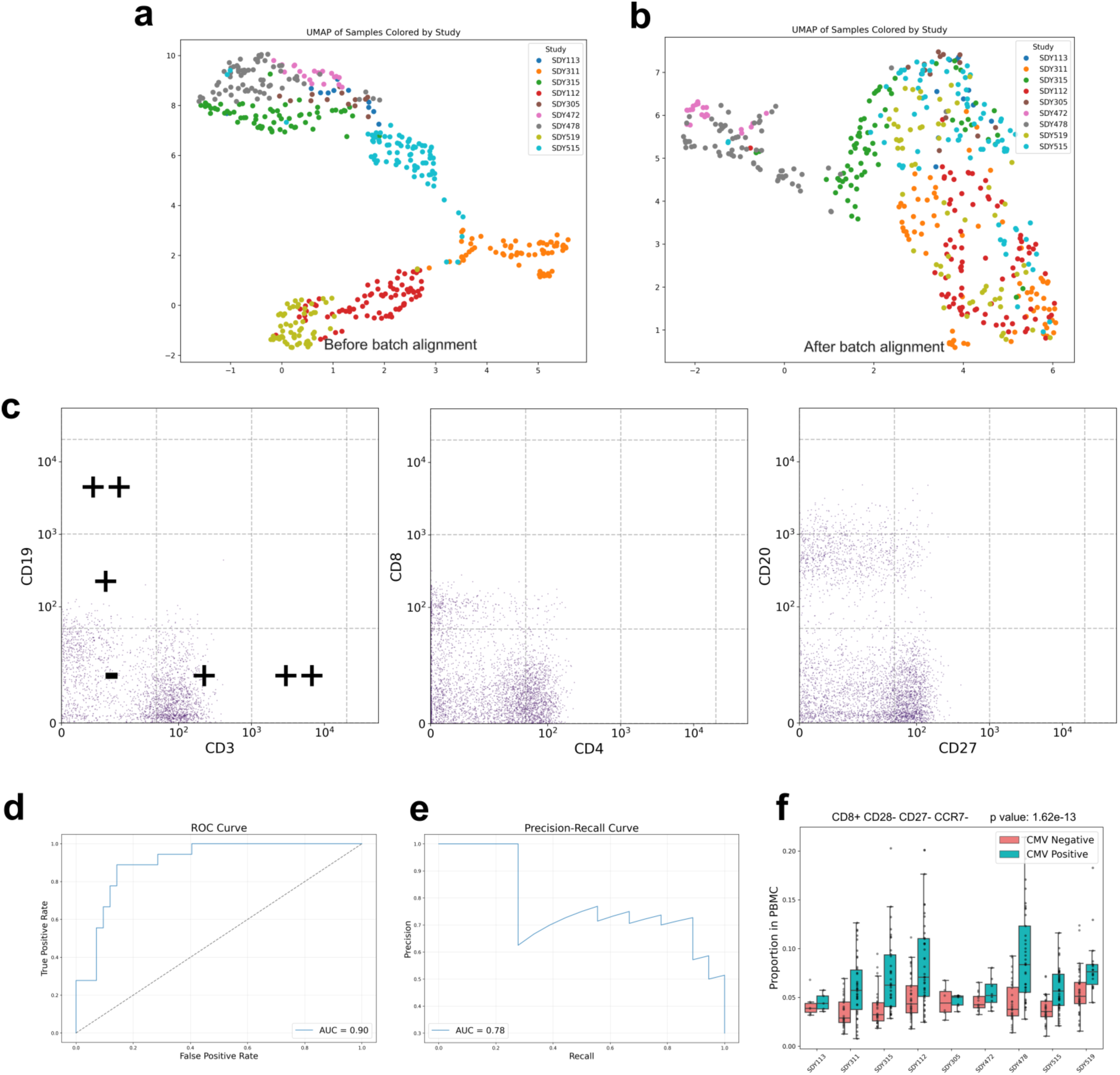
Batch alignment and classification analysis of cytometry data for CMV status prediction. (a) UMAP visualization of samples coloured by study batch before batch alignment, showing clear batch-specific clustering. (b) UMAP visualization after batch alignment, demonstrating improved mixing of samples across batches. (c) Representative flow cytometry scatter plots showing population distributions and voxel-gating intervals for key markers (CD3, CD19, CD4, CD8, CD27, CD20). (d) Receiver Operating Characteristic (ROC) curve for CMV status classification showing an AUC of 0.90. (e) Precision-Recall curve for the classification model with AUC of 0.78. (f) Box plots showing the proportion of CD8^+^/CD28^-^/CD27^-^/CCR7^-^ T cells across different study batches, stratified by CMV status.

Our re-analysis provides deeper phenotypic resolution than the original work and identified several highly significant CD8^+^ T cell populations that share key features with the original findings. The most significant hyper-voxel combination (p = 1.62e^-13^) identified a cell subset with CD8^+^ CD28^-^ CD27^-^ CCR7^-^ which is concordant with the original work, as shown in the box plots (Figure 5f). These plots clearly demonstrate the increased proportion of this specific T cell phenotype across different study batches in CMV-positive individuals compared to CMV-negative individuals. Notably, while Hu et al. emphasized CD94 expression, our analysis highlights the importance of CD28 and CCR7 negativity, markers typically associated with terminal differentiation and effector memory phenotypes. The subsequent combinations maintain CD8^+^/CD27^-^/CCR7^-^ as a core signature while varying other markers (CD127, CD4, TCRGD, CD45RA), suggesting these cells represent a robust CMV-associated population. The consistent minimum fraction (0.008) and similar maximum fractions (0.214-0.218) across most combinations suggest these may be overlapping populations. The inclusion of CD45RA in the fifth combination (CD8^+^/CD45RA^-^/CD27^-^/CCR7^-^) particularly supports the conclusions of the original work^24^ that these cells span multiple differentiation states, including effector memory phenotypes.

**Figure 5.**
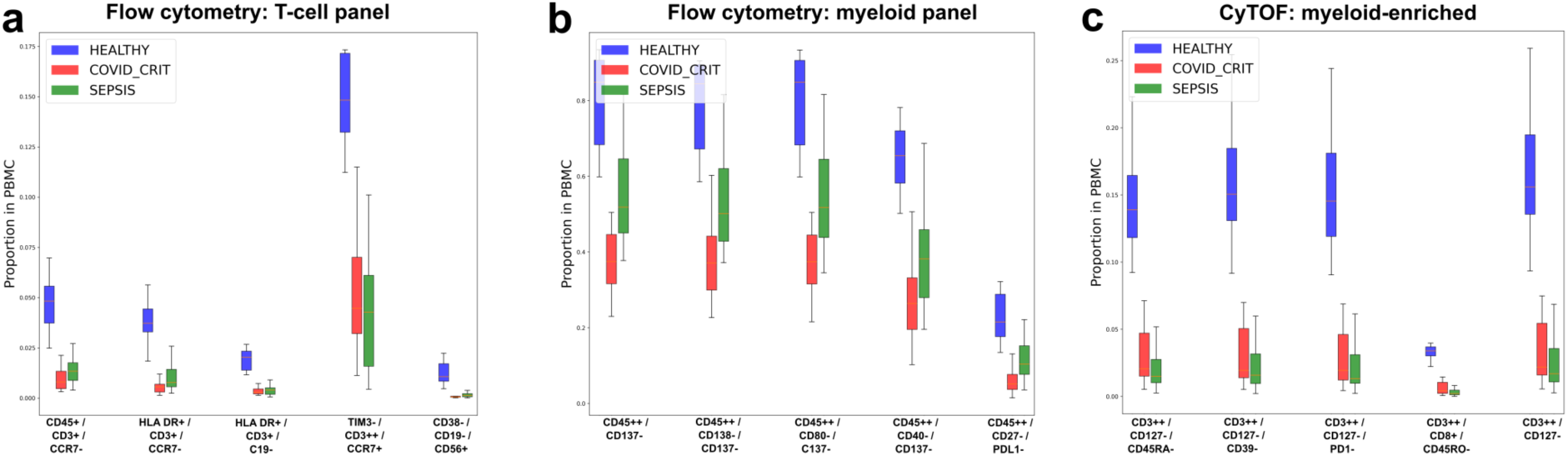
Comparative analysis of immune cell populations in healthy controls, COVID-19 critical patients, and sepsis patients across three different analytical platforms. **(a)** Flow cytometry analysis of T-cell populations. **(b)** Flow cytometry analysis of myeloid cell populations. **(c)** Mass cytometry (CyTOF) analysis of myeloid-enriched cell populations Box plots show the proportion of cells in peripheral blood mononuclear cells (PBMCs) for each marker combination.

### Discrimination between sepsis and COVID-19 patients using CyTOF and multicolour flow cytometry

As a further test with real-world data, we partially re-analysed the cytometry dataset generated by the COMBAT consortium^25,26^ from individuals infected with COVID-19. This dataset presents significant analytical challenges typical of clinical cytometry data: high patient-to-patient variability, complex disease states, and technical variation across multiple collection sites. We re-analysed their cytometry dataset using our VoxelCoder pipeline, demonstrating that automated multi-dimensional analysis can extract additional immunological insights and achieve robust disease classification.

The flow cytometry measurements were made across 6 experimental batches from individuals clinically classified with mild, severe and critical COVID-19 infection. The dataset also contains flow samples from patients with sepsis and flu, as well as healthy control samples (see Supp. Info. Table 2). To simplify the analysis task, we only focus on the following sample classes: 1. health, 2. critical COVID infection, 3. sepsis. Before sample classification and biomarker discovery analysis were made, we apply our autoencoder batch alignment model to the entire dataset.

In a hyper-voxel framework using VoxelCoder, we developed a classification model to predict the three infection-status classes. The classification model achieved strong performance in distinguishing between patient groups, with area under the ROC curve values of 0.99, 0.90, and 0.92 for healthy controls, COVID-critical, and sepsis patients, respectively (see Supplementary Figure 2). Analysis of classification accuracy revealed perfect discrimination of healthy controls (100% correct classification), while COVID-critical and sepsis cases showed some degree of overlap, with 78.6% correct classification of COVID-critical cases and 83.3% correct classification of sepsis cases. This overlap may reflect shared immunological features between these acute inflammatory conditions.

Our analysis (see Figure 5 a,b,c) revealed distinct T cell populations with discriminatory power between COVID-19, sepsis, and healthy controls. Most notably, we identified a CD3^++^/CD45RA^++^/CD8^+^ population, likely representing naive cytotoxic T cells, and several CD4+ subsets with varying expression of activation markers (CD38, CD25) and migration markers (CCR7). The CD4^+^/CD45RA^++^/CCR7^++^ signature suggests involvement of naive helper T cells, while CD4^+^ populations expressing high levels of CD38 and CD25 indicate increased T cell activation states. These findings highlight the differential immune responses in COVID-19 versus sepsis, particularly in the naive and activation status of both CD4^+^ and CD8^+^ T cell populations (Figure 5).

## Discussion

We show here that batch alignment of cytometry data with an autoencoder neural network architecture achieves sufficiently normalised marker intensity signals to enable use of a multi-dimensional representation of this information in subsequent machine learning classification tasks. This ‘hyper-voxel’ representation does not involve dimensionality reduction, remains fully interpretable and establishes of a new conceptual framework for working with large, integrated cytometry datasets for sample classification and predictive model building tasks. Furthermore, the method eliminates the requirement for per-plate normal controls and manual gating, potentially increasing practical utility and analytical reproducibility. This provides a simplified path towards to automated analysis of very large cytometry datasets that are both longitudinal and multi-centre.

A critical prerequisite for application of the hyper-voxel methodology described here is the accurate alignment of samples to remove technical batch effects that are ubiquitous in cytometry data. The method presented here incorporates batch alignment as an integral first step prior to downstream classification task. In this way it differs from many current machine learning approaches, including deep neural networks, that commonly struggle with prospective applications due to batch effects that are significantly outside the original model training distribution. By optimizing both reconstruction accuracy and distribution matching through a combined loss function, this normalization step aims to ensure that cells expressing similar marker levels will consistently fall within the same voxels, regardless of their batch of origin. The VoxelCoder approach integrates both batch alignment and the hyper-voxel representation of cell counts, and hence mitigates technical variations in future data acquisition, a key consideration in prospective clinical applications.

We benchmarked our autoencoder-based batch alignment approach using a controlled murine dataset with synthetically introduced batch effects, demonstrating that VoxelCoder effectively addresses technical and batch-related variation in cytometry data. Critically, VoxelCoder retains more biological signal than popular methods, including CytoNorm^9,10^ and Harmony^13^, as evidenced by superior Area Under the Curve (AUC) values and statistical significance of identified features in real experimental datasets. This performance highlights the critical dual requirement in batch correction: minimizing technical variation while simultaneously preserving biological signal. VoxelCoder’s distribution-matching approach achieves this balance through an autoencoder neural network that learns dataset-specific transformations, removing technical artifacts while retaining biologically meaningful variations. The autoencoder framework’s ability to handle non-linear batch effects while maintaining population structure preserves the resolution of input data, enabling robust analysis of large-scale studies. This strength is best demonstrated in downstream applications, including two challenging clinical datasets: predicting CMV serostatus from cytometry data spanning nine independent studies, and discriminating between COVID-19 and sepsis patients in the COMBAT study across six experimental batches.

Our approach achieves classification performance comparable to black-box machine learning methods on input flow cytometry and CyTOF datasets that were not considered valuable for classification tasks by the original projects that generated them. Unlike black-box machine learning classifiers, our multi-dimensional representation of normalised data maintains complete interpretability of cellular counts and marker intensity features. This represents an important advantage over deep learning approaches that require complex post-hoc analysis. The hyper-voxel analysis framework complements rather than replaces manual gating, systematically exploring the multi-dimensional marker space to identify significant populations for validation through conventional gating strategies, while enabling comprehensive examination of marker combinations that might be overlooked in manual analysis. Finally, unlike existing methods that require common controls across batches, our method achieves normalization by learning marker distributions from a reference batch, making it applicable to studies where matched samples are unavailable. This capability is particularly valuable for integration of public datasets, where matched controls are often not available across different studies.

For this work we generated a benchmarking dataset for the evaluation of the performance of batch alignment methodology. We have applied this here to benchmark the performance of our tool, VoxelCoder, with three other prominent batch normalisation tools: CytoNorm^9,10^, CyCombine^12^ and Harmony^13^. As the cells in each technical replicate for each mouse were derived from the same original splenocyte suspension, a perfect batch alignment would result in little or no difference between technical replicates. We find that VoxelCoder has equivalent performance with CyCombine and Harmony for this task, though we have not attempted to quantitate experimental variation or sampling error in splitting of the original cell suspensions. Furthermore, a gender bias between the biological replicate mice appeared to skew some batch alignment results. Taking advantage to subtle cellular phenotype differences identified between male and female mice^27^ we observed that VoxelCoder removed less true biological signal during batch alignment than other tools, especially Harmony. This represents an important facet to cytometry batch normalisation, especially when applying this to detection of subtle cellular phenotypes of human diseases.

When applied to a CMV serostatus prediction task, the VoxelCoder method identified populations consistent with previous findings^24,28^ while providing additional phenotypic resolution. A highly predictive cell population of CD8^+^/CD28^-^/CD27^-^/CCR7^-^ T cells identified by the hyper-dimensional representation of this data aligns with known CMV-associated effector memory populations^24^. Similarly, in the COVID-19/sepsis classification task, the method revealed distinctive T cell activation signatures that differentiated between disease states while maintaining direct interpretability. The original study^25^ did not identify these signatures, as shown by their reported discriminative features.

Future work could explore two new directions. First, the development of adaptive thresholding approaches for marker discretization could improve robustness across different experimental contexts. Second, systematic validation studies comparing our voxel-based populations against published manual gating strategies using open-access datasets could help quantify the correspondence between these approaches. Specifically, analyzing how frequently cells within established manual gates map to specific hypervoxels could provide valuable insights into the relationship between these different analysis paradigms.

We show that batch alignment with autoencoders allows a simple representation that greatly improves classifier performance. Through effective batch alignment, larger cytometry data may be now integrated and interpreted without significant loss of original data resolution. This greatly enables data science approaches to using this modality for longitudinal studies of individuals, multicentre studies of cohorts and any large cytometry study conducted over significant timescales with potentially heterogeneous cytometry equipment. This will also allow better integration of datasets from public repositories of flow- and CyTOF mass cytometry datasets.

## Methods

### Generation of benchmark mouse synthetic batch-effect flow cytometry dataset

A benchmarking dataset was produced from flow cytometry of three biological replicate wild-type C57BL/6 mice (two female, one male) and two mutant C57BL/6 strains (*Kenobi*, a CD79a null strain^29^) and a RAG-1 null strain^30^). The dataset was generated from splenic cell suspensions from each mouse split to three technical replicates and prepared such as to introduce a controlled, synthetic batch effect from the same input cells. The synthetic batch effect was produced through minor manipulation of antibody dilutions in three marker panels (see below; Table 4).

**Table 3.** Top 5 most statistically significant voxels by CMV status discrimination. Calculated with VoxelCoder aligned dataset.

| Feature | p-value | Min. occupancy | Max. occupancy |
| --- | --- | --- | --- |
| CD8<sup>+</sup> / CD28<sup>-</sup> / CD27<sup>-</sup> / CCR7<sup>-</sup> | 9.89e-17 | 0.008 | 0.214 |
| CD8<sup>+</sup> / CD27<sup>-</sup> / CD127<sup>-</sup> / CCR7 | 1.27e-16 | 0.080 | 0.215 |
| CD8<sup>+</sup> / CD4<sup>-</sup> / CD27<sup>-</sup> / CCR7<sup>-</sup> | 1.60e-16 | 0.082 | 0.215 |
| TCRGD<sup>-</sup> / CD8<sup>+</sup> / CD27<sup>-</sup> / CCR7<sup>-</sup> | 1.81e-16 | 0.084 | 0.218 |
| CD8<sup>+</sup> / CD27<sup>-</sup> / CCR7<sup>-</sup> | 2.16e-16 | 0.084 | 0.218 |

**Table 4.** Details of antibodies and dilutions used for each panel in the mouse synthetic batch effect dataset. Each antibody is described by the marker to which it binds, the attached fluorophore, the dilution factor in each panel and the catalogue and supplier of the antibody. Panel 1 is the optimal antibody concentrations determined by antibody titration, with minor changes in the further two panels to induce a synthetic batch effect.

| Marker | Fluorophore | Panel 1<br>dilution<br>factor | Panel 2<br>dilution<br>factor | Panel 3<br>dilution<br>factor | Supplier | Catalogue No. |
| --- | --- | --- | --- | --- | --- | --- |
| CD3 | FITC | 200 | 200 | 300 | BioLegend | 100204 |
| CD25 | PE | 400 | 400 | 400 | BioLegend | 102008 |
| CD8 | CD8 | 100 | 400 | 400 | BD | 563786 |
| CD44 | PacBlue | 100 | 400 | 400 | BioLegend | 103020 |
| CD62L | BV605 | 400 | 600 | 600 | BioLegend | 104438 |
| CD19 | BV510 | 600 | 800 | 800 | BioLegend | 115546 |
| IgD | PerCPCy5.5 | 200 | 800 | 800 | BD | 564273 |
| IgM | R718 | 100 | 100 | 600 | BD | 752171 |
| NK1.1 | APC | 200 | 400 | 600 | BD | 550627 |
| Ly6C | PECy7 | 200 | 200 | 400 | BioLegend | 128018 |
| B220 | BUV737 | 300 | 600 | 800 | BD | 612838 |
| CD4 | AF700 | 200 | 300 | 600 | BioLegend | 100430 |
| LiveDead | ef780 | 400 | 600 | 800 | ThermoFisher | 65-0865-14 |

Mouse spleens were collected at into 3 mL FACS buffer (2.5% FBS, 0.1% sodium azide, 0.01% EDTA, 10% PBS) and mashed through a 70 µm cell strainer. Cells were transferred into 15 mL falcon tubes, centrifuged at 465 xg for 5 min at 4°C, following which the supernatant was discarded. Red blood cells were lysed by resuspending the pellet in 3 mL 1X lysis buffer (ThermoFisher Scientific, 00-4300-54), incubated for 1 min at room temperature. Post incubation, lysis buffer was diluted by the addition of 7 mL of FACS buffer and centrifuged at 465 g for 5 min at 4°C. Supernatant was subsequently discarded and the cells washed by resuspending in 10 mL FACS buffer before being centrifuged again at 465 xg for 5 min at 4°C. The cell pellet was resuspended in 0.5 mL FACS buffer and the total cell count calculated using the Luna-II Automated Cell Counter (ThermoFisher Scientific). Cells were plated into a 96 well round bottom plate.

Cells were blocked in 25 µl of 2X Fc block (BD, Ms CD16/CD32 Pure 2.4G2, 553142) for 5 min before the addition of 25 µl 2X Live Dead stain (ThermoFisher Scientific, Fixable viability dye E780, 65-0865-14). After staining, cells were washed in 200 µl 1xPBS and centrifuged at 465 xg for 5 min at 4°C. For each biological replicate mouse, the total cell suspensions were divided into three equal parts. Cells were stained with 50 µl of antibody cocktail (in brilliant stain buffer BD Biosciences, 566349) with panels diluted to different antibody concentrations (Table 4) or the respective single colour control at 4°C for 30 min before washing in FACS buffer. Cells were fixed using eBioscience Fix buffer 00-5523-00 according to manufacturer’s instructions. Post fixing, cells were washed twice with FACS buffer before resuspending in 80 µl of FACS buffer for acquisition on the LSRFortessa X-20 (BD).

### Dataset pre-processing

Flow cytometry data was processed using a custom Python function built on the FlowKit framework^31^. The function accepts Flow Cytometry Standard (*.fcs) files and performs sequential preprocessing steps: First, compensation is applied to correct for fluorescence spillover between channels. Cellular debris and non-single-cell events were then removed using sequential polygonal gates defined in the forward and side scatter dimensions. For flow cytometry data, a logicle (biexponential) transformation with parameters T=262144, W=0.5, M=4.5, and A=0 was applied, while mass cytometry (CyTOF) data was transformed using arcsinh transformation with a cofactor of 5 followed by scaling by a factor of 1/8. These transformations, which are standard practice in cytometry analysis, provide appropriate scaling for both negative and positive values while maintaining resolution of low-intensity signals. For training the autoencoder batch alignment model, a fixed count of 30,000 cells was subsampled from each sample in the reference batch.

CMV data sourced from repository by Hu et al^32^, and COMBAT COVID-19 data sourced from public repository made available by study authors^26^.

### Batch alignment via deep neural network autoencoder

The batch alignment method was implemented as a custom PyTorch ^33^ model using an autoencoder neural network architecture. The network consisted of three encoding layers (64, 32, and 16 nodes) and three symmetrical decoding layers, with ReLU activation functions between layers. The autoencoder was trained to transform input data from different batches to match a reference distribution while preserving biological signal. One batch was randomly selected as the reference, and the model was trained using a combination of mean squared error (MSE) loss and a custom histogram loss function that encouraged the transformed data to maintain similar marker distribution shapes as the reference batch.

The histogram loss term was computed by comparing normalized histogram distributions between the network output and reference data across all markers, using Wasserstein distance as the similarity metric. This dual loss function helped ensure that both individual cell measurements and overall population distributions were preserved during batch alignment. The model was trained for 1,200 epochs using the Adam optimizer^34^ with a learning rate of 0.002 and a batch size of 1,024 cells. The histogram loss term was weighted by a factor β=0.002, which was gradually decreased during training to allow fine-tuning of individual cell measurements in later epochs while maintaining population-level distribution matching.

### Voxel-gating for multi-marker population analysis

Following batch alignment, we implemented a systematic voxel-based analysis strategy to characterize cellular populations. For each marker, expression values were discretized into three levels (LOW, MED, HIGH) using fixed intensity thresholds (<0.3, 0.3-0.6, >0.6 in normalized intensity units). We then generated all possible two- and three-marker combinations from the panel, considering only biologically relevant marker combinations (Supp. Info. Table 1) within each lineage. This strategy was necessary to mitigate the extreme sparsity and computational overhead of the full and large multi-dimensional space, through considering a subset of marker combinations with biological relevance (e.g., T cell markers were only combined with other T cell markers). For each marker combination, we computed the proportion of cells falling within each possible discretized state combination, creating a high-dimensional feature vector of population frequencies. For example, a combination of three markers would generate 27 possible states (3³ combinations of LOW/MED/HIGH), with the frequency of cells in each state serving as a feature for downstream analysis.

These population frequency features were compiled into a sample-by-feature matrix where each row represented a sample, and each column represented the frequency of cells in a particular marker combination state. This matrix served as input for subsequent machine learning classification tasks, providing a comprehensive yet interpretable representation of the cellular composition of each sample. This approach effectively transforms complex single-cell data into discrete population-based features while maintaining the ability to examine high-dimensional marker relationships.

### Classification and identification of significant features

Following voxel-based feature generation, we implemented machine learning classification approaches tailored to each dataset. For the CMV dataset^24,28^, we employed a hold-out validation strategy, designating a specific batch (SDY519) as the test set while training on all remaining batches. For the COVID-19 dataset^25,26^, we implemented batch-wise cross-validation, where the model was trained on n-1 batches and tested on the remaining batch. In both cases, we used XGBoost classifiers^35^ to distinguish between patient groups (CMV-positive vs. CMV-negative for the first dataset; healthy controls, COVID-19 critical patients, and sepsis patients for the second) using population frequencies derived from our voxel-gating analysis. Features were standardized using z-score normalization, and class weights were applied to address class imbalance. The models were optimized using 500 trees with a maximum depth of 6, and thresholds for prediction were fine-tuned using F1 scores.

Significant features were identified through a combination of statistical testing and machine learning importance metrics: initial feature selection was performed using one-way ANOVA to identify marker combinations that showed significant differences between patient groups.

## Code Availability

An implementation of this strategy, VoxelCoder, accompanies this work, accessible at: http://voxelcoder.cloud

## Acknowledgments

The authors thank the National Computational Infrastructure (Australia) for continued access to significant computation resources, and the Cytometry, Histology and Advanced Spatial Multiomics Facility at the John Curtin School of Medical Research.

## SUPPLEMENTARY INFORMATION

**Supplementary Figure 1.**
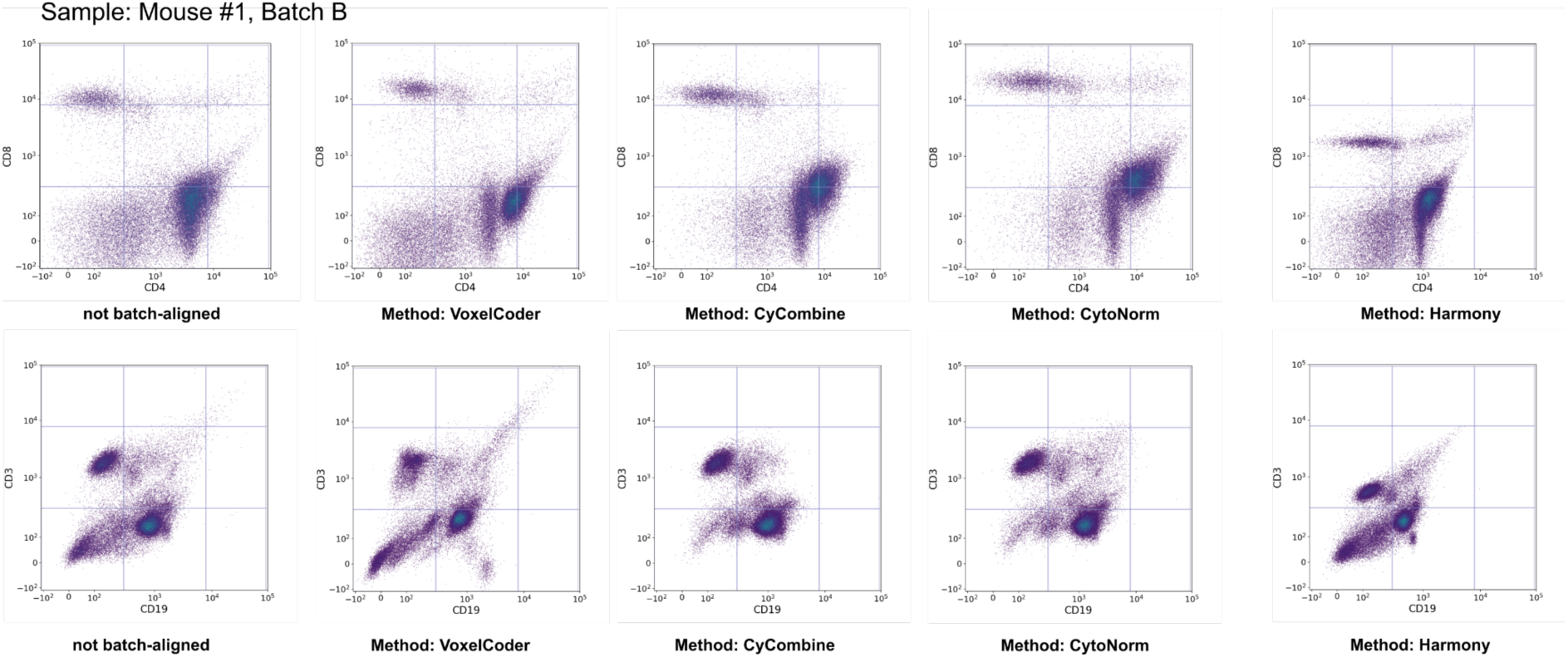
Scatter plots showing the performance of four different batch normalization methods (VoxelCoder, CyCombine, CytoNorm, and Harmony) alongside non-batch-aligned controls for Mouse #1, Batch B. The top row shows a scatter plot of CD8 vs. CD4, while the lower row shows CM19 vs. CD3.

**Supplementary Figure 2:**
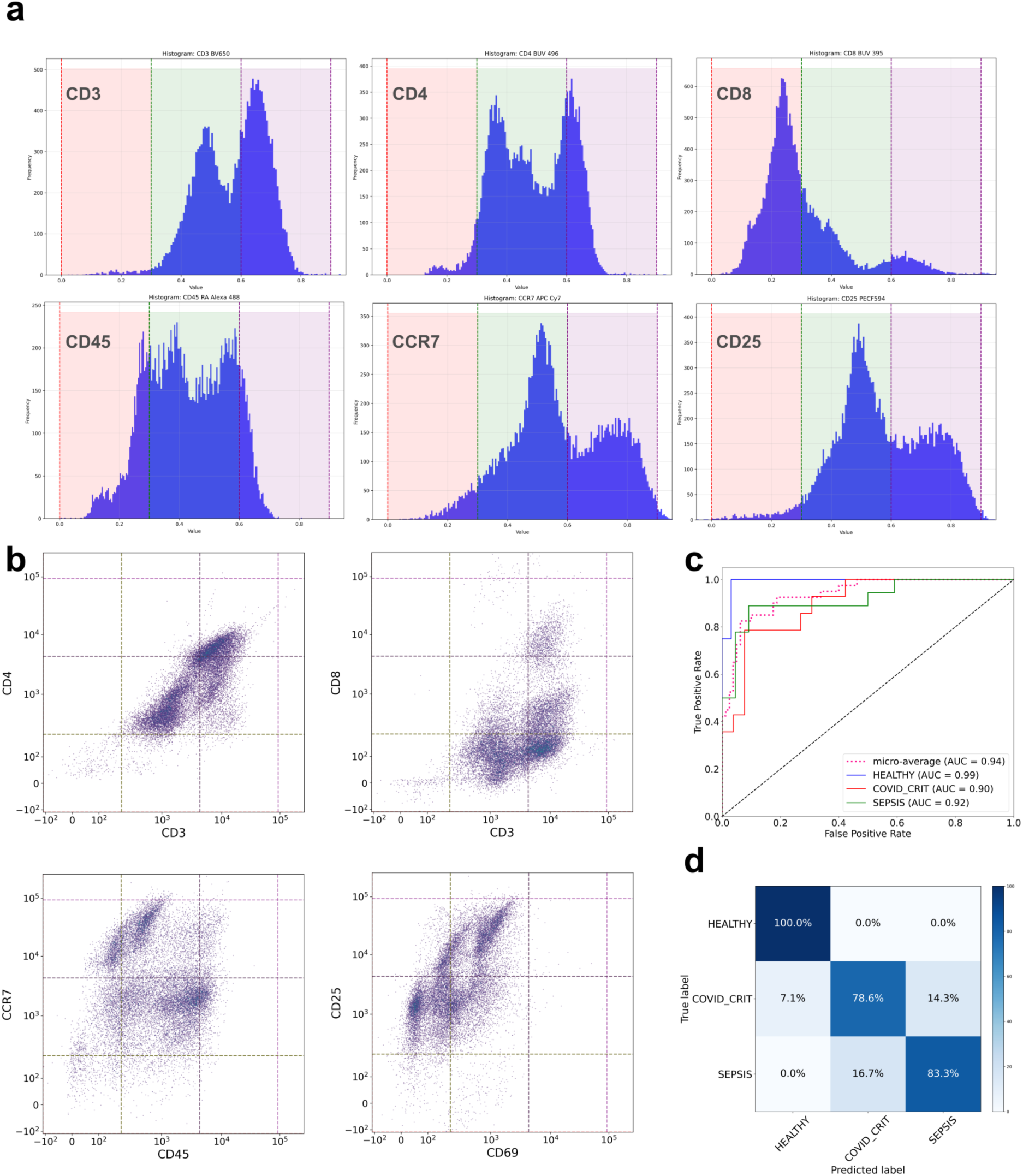
Flow cytometry analysis and machine learning classification of immune cell populations across patient groups in COMBAT dataset (T-cell panel). **(a)** Histogram distributions of key T-cell surface markers (CD3, CD4, CD8) and activation/migration markers (CD45, CCR7, CD25), showing expression patterns with negative (pink), intermediate (green), and positive (purple) gating intervals. **(b)** Scatter plots demonstrating the co-expression relationships between paired markers, including T-cell defining markers (CD3 vs CD4, CD3 vs CD8) and activation markers (CD45 vs CCR7, CD69 vs CD25). **(c)** Receiver Operating Characteristic (ROC) curves showing the classification performance for distinguishing between healthy controls, COVID-19 critical patients, and sepsis patients, with area under the curve (AUC) values indicated for each group. **(d)** Confusion matrix displaying the classification accuracy of the model across the three patient groups, with percentages indicating correct and incorrect classifications.

**Supplementary Figure 3.**
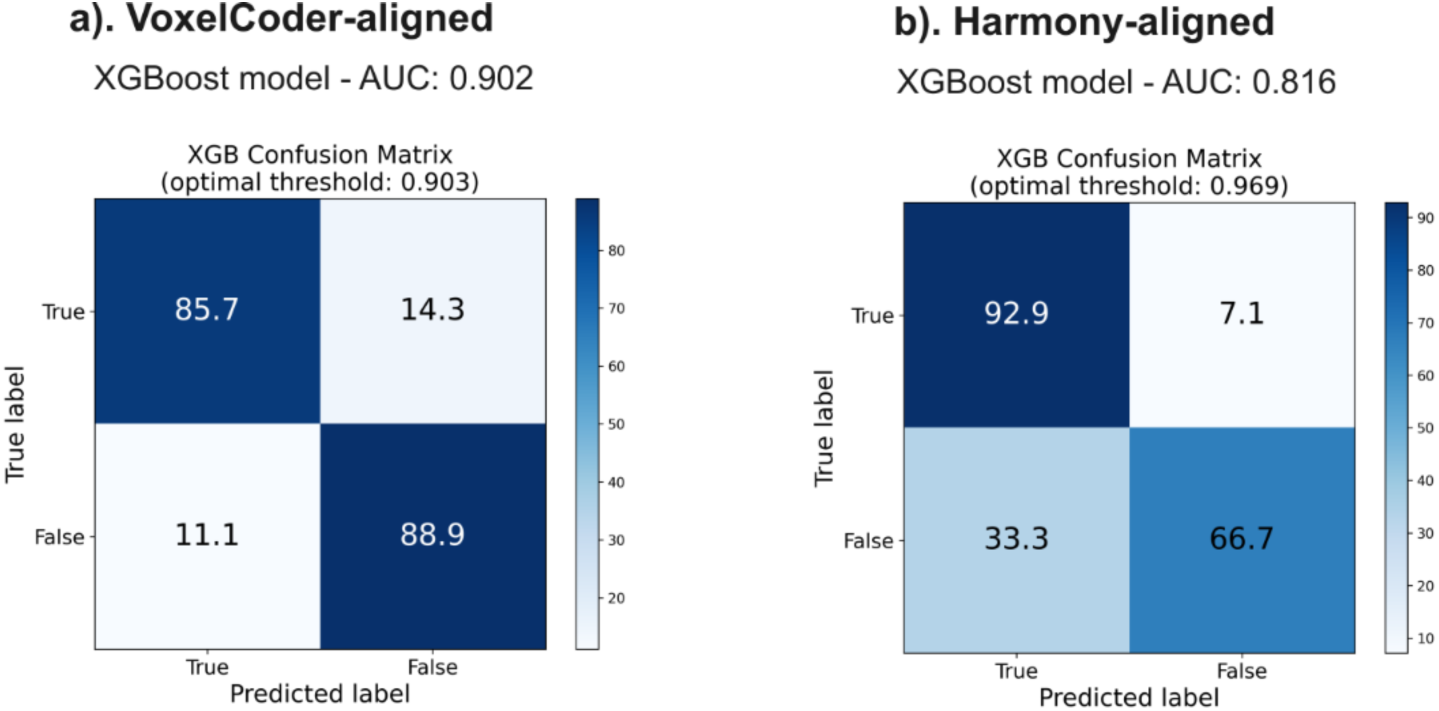
XGBoost classifier performance on CMV-status discrimination task, comparing a). VoxelCoder-aligned dataset vs. b). Harmony-aligned dataset.

## Supplementary Tables

**Supplementary Table 1.** Markers used for immune cell population characterization in CMV study. Mass cytometry markers are grouped by major immune cell lineages (T cells, B cells, NK cells, and myeloid cells), showing the lineage-specific combinations used for multi-dimensional population analysis in the voxel-based framework.

|  |  |
| --- | --- |
| T cell markers | 'TCRGD', 'CD8', 'CD45RA', 'CD4', 'CD3', 'CD28', 'CD27', 'CD127', 'CCR7' |
| B cell markers | 'CD19', 'CD20', 'CD24', 'IGD', 'CD27' |
| NK cell markers | 'CD56', 'CD16', 'CD94', 'CD161' |
| Myeloid cell markers | 'CD14', 'CD33', 'HLADR' |

**Supplementary Table 2.** Distribution of patient samples in entire COMBAT dataset across disease categories and control groups.

| Classification | Sample count |
| --- | --- |
| COVID (severe) | 40 |
| Sepsis | 22 |
| COVID (mild) | 18 |
| COVID (critical) | 18 |
| COVID (health care worker) | 13 |
| Flu | 11 |
| Healthy volunteer | 10 |
| LDN treated | 2 |
| **Total** | **134** |

